# Assessment of trace metals and arsenic contamination in water and sediments in a former mining area of the river basin after an ecological disaster, Slaná River, Slovakia

**DOI:** 10.64898/2026.10.08.757544

**Authors:** Marek Helczman, Marian Tomka, Anton Kovacik

## Abstract

This study assessed the spatial distribution of trace metals and arsenic in the water and sediment of the Slaná River (Slovakia) approximately one and a half years after the 2022 release of mine water from the former Nižná Slaná siderite mine. Samples were collected in September 2023 at four river sites from Gočovo to Lenártovce (L1–L4) and at five points within 50 m of the mine-water discharge (S0–S4), and the data were evaluated using non-parametric tests, principal component analysis, contamination factors (CF) and the pollution load index (PLI). At the discharge point, the water contained 149 mg/L Fe, 27 mg/L Mn and 0.29 mg/L As; beyond this point, however, most metals in the filtered river water were below the limit of detection, and only sulphur remained clearly elevated downstream (35.7 mg/L at L2 vs. 6.4 mg/L at the reference site). In contrast, the sediment retained the contamination: As reached 7920 mg/kg and Cd 39 mg/kg at the discharge, exceeding the probable effect concentrations of freshwater sediment quality guidelines more than 200-fold and almost eightfold, respectively, and As remained above this level at the first downstream site (47 mg/kg). While As, Hg, Mn and Ni declined rapidly downstream, S and Al accumulated toward the most distant site, so that the PLI remained uniform along the river (1.06–1.16; p = 0.246). Despite the apparent recovery of the water column, the sediments of the Slaná River thus remain a persistent reservoir of arsenic and trace metals and a potential source of chronic exposure for biota; repeated sampling across seasons is needed to confirm this pattern.

## 1. Introduction

Mining is one of the most persistent sources of anthropogenic stress on river ecosystems, and its effects persist in water systems for a long time after mine closure (Bensadi C Azzoug, 2026; Biedunkova C Kuznietsov, 2024; Macklin et al., 2023). Acid mine drainage (AMD) affects the ecosystem simultaneously at several levels—it alters hydrochemical conditions, the mineralogical composition of water and sediments (Favas et al., 2024; Xia et al., 2017), the structure of microbial and periphyton communities (Luís et al., 2021), and fish trophic networks (Kovacik et al., 2025; Krause et al., 2019). This broad-spectrum impact is crucial for ecosystem recovery, as improved water quality alone may not lead to the restoration of a functional ecosystem. As long as abiotic barriers persist, the system may remain in an altered but stable state (Ghosh C Ghoshal, 2026; Suding et al., 2004). From a biogeochemical perspective, the long-term persistence of heavy metals and metalloids in such systems is based on their accumulation in sediments (Biedunkova C Kuznietsov, 2025; Khan et al., 2026; Uddin et al., 2022). The key mechanism is the oxidation of Fe(II) and its subsequent coprecipitation with Fe(III) oxyhydroxides and hydrosulphates (Thomas et al., 2018). Sediment therefore acts as a chronic reservoir (De Almeida Ribeiro Carvalho et al., 2022; Shen et al., 2025) that regulates the bioavailability of trace metals for future generations of biotic communities and maintains an exposure pathway even when the water column is showing signs of recovery (Morariu et al., 2025). Periphyton biofilms function in these systems as the entrance gate for metals into the food web, accumulating elevated concentrations from the aqueous phase and mediating trophic transfer to fish (Bonnineau et al., 2020; Wen et al., 2024). Bioaccumulation of toxic elements in the target organs of fish has been repeatedly documented in aquatic ecosystems affected by mining (Bisht et al., 2026; Kovacik C Helczman, 2025; Ujan et al., 2025).

In Central Europe, the Spiš-Gemer mining region is one of the areas with the longest history of intensive mining, with a long history of mining of ferrous siderite enriched with Mn and associated sulphides containing trace elements such as As, Pb, Zn, and Hg (Demková et al., 2019, 2020; Fazekašová C Fazekaš, 2020). In February 2022, a long-accumulated, highly mineralized mine water actively leaked through the drainage system directly into the Slaná River. The river immediately turned ochre-brown/orange in color, and this was followed by a drastic degradation of fish communities over tens of kilometers downstream (Fenyo C Than, 2022; King, 2022; Tanács, 2022). Despite media and political attention, the condition of the Slaná River remains poorly described in the scientific literature. Following this ecological incident, the installation of filters to prevent the continued leakage of pollutants into the watercourse was confirmed. A slow recovery of the ecosystem to its original state is expected in time.

The aim of this study is therefore to provide a detailed abiotic profile of the Slaná River approximately 1.5 years after the events, which will serve as an environmental framework for the parallel biological assessment of fish communities. In this study, we characterize the micro-gradient of contamination in the immediate zone of the mine discharge using five sampling points (50 and 20 m upstream of the discharge, directly at the discharge, 20 and 50 m downstream of the disposal point) and quantify the spatial dynamics of metal and trace element attenuation in surface water and sediment along the river gradient from Gočovo through Henckovce and Plešivec to the final sampling point in the Slovak Republic at Lenártovce. Using multivariate analysis and integrated load indices (CF, PLI), we assess the residual ecological risk of the sedimentary component as a potential source of chronic exposure for biota. The present analysis is conceptually designed to link to the biological indicators in the ongoing study of fish communities in the Slaná River and to provide a geochemical context for interpreting the observed ecotoxicological effects.

## 2. Material and Methods

### 2.1 Location and sample collection

In September 2023, we collected water and sediment at five river sites: Gočovo (L1, upstream reference site; 48°45’31.3”N 20°24’53.6”E), Nižná Slaná – mine-water discharge point (S0-S4; 48°44’05.6”N 20°24’58.1”E), Henckovce (L2; 48°42’51.3”N 20°26’06.9”E), Plešivec (L3; 48°32’46.9”N 20°23’46.5”E)and Lenártovce(L4; 48°18’14.9”N 20°18’51.8”E), along 100-m river reaches, taking samples every 5-10 m across each reach. At each sampling point, approximately 500 mL of river water for trace-metal analysis and about 100 g of surficial bottom sediment (0–5 cm) were collected, in line with recommended sample volumes and masses for metals monitoring in freshwaters and river sediments. In the immediate impact zone of the pollution source, additional samples were taken directly at the mine-water outlet (S0), 20 m downstream (S1), 50 m downstream (S2), as well as 20 m (S3) and 50 m (S4) upstream of the discharge. The spatial layout of all sampling locations and river reaches is shown in Figure 1.

**Figure 1.**
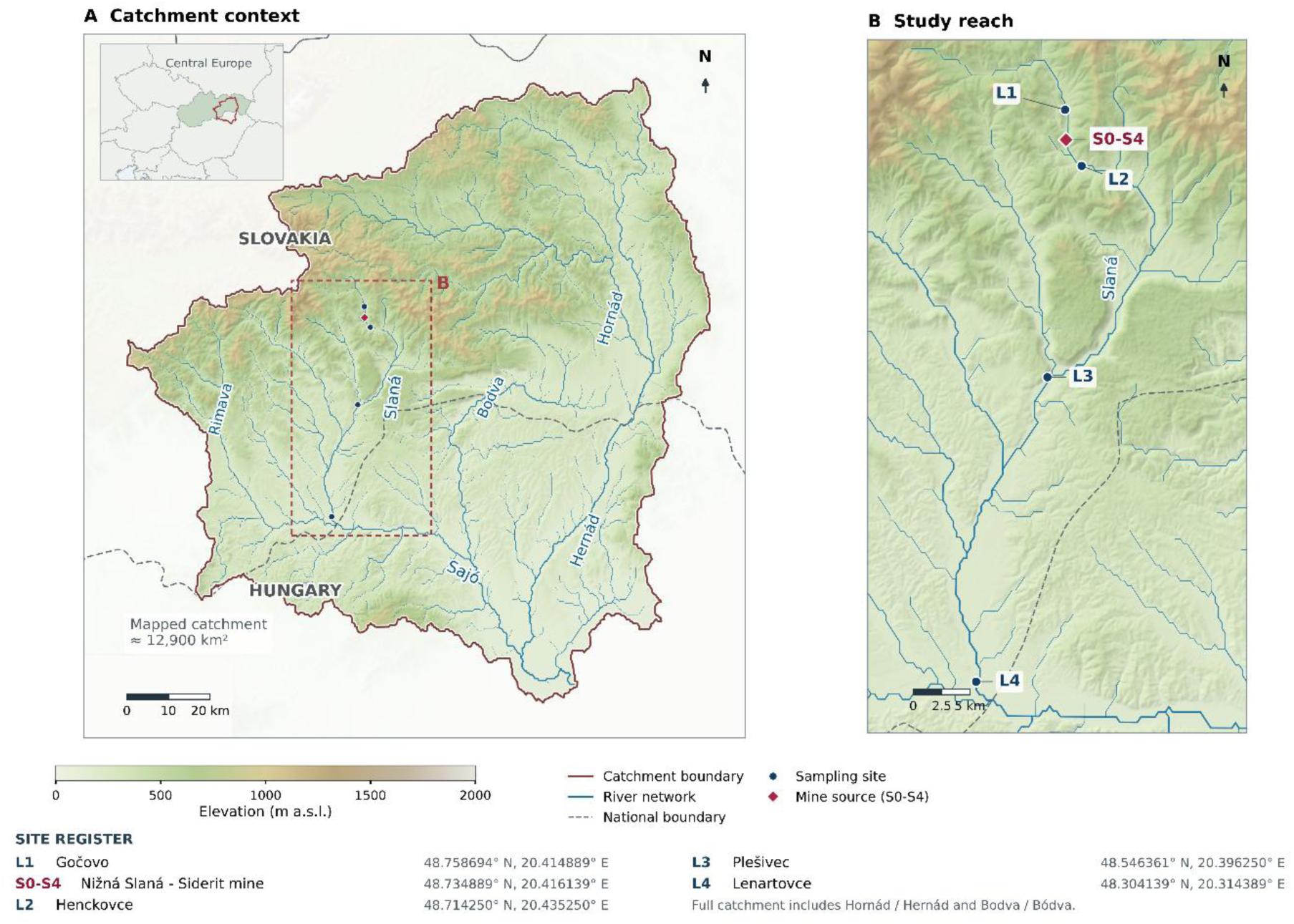
Topographic and hydrological setting of the Slaná River study area. **(A)** The transboundary Slaná/Sajó catchment, including the Hornád/Hernád and Bodva/Bódva tributary catchments, with an inset showing its location within Central Europe. The dashed red rectangle indicates the area enlarged in panel B. **(B)** The investigated Slovak river reach, showing the contamination source at the Siderit mine in Nižná Slaná (S0 -S4) and the water and sediment sampling sites at Gočovo (L1), Henckovce (L2), Plešivec (L3), and Lenartovce (L4). Elevation and shaded relief were derived from Mapzen Terrain Tiles; the river network and catchment boundary were obtained from HydroRIVERS v1 and HydroBASINS v1c, respectively. National boundaries were obtained from Natural Earth. Maps are displayed in WGS 84 / UTM zone 34N (EPSG:32634).

### 2.2 Sample preparation for trace metals analysis (ICP-OES)

Dried (finely homogenised) river sediment samples were prepared for ICP-OES analysis using a closed-vessel, microwave-assisted acid digestion system speedwave XPERT (Berghof Products + Instruments GmbH, Eningen, Germany). For sediment, up to 0.5 g of dry material was weighed into digestion vessels and a mixture of 8 mL concentrated HNO₃ (TraceSELECT®, Honeywell Fluka, Morris Plains, USA) and 2 mL H₂O₂ (30%, for trace analysis, Merck Suprapur®) was added. After microwave digestion (15 min heating to 200 °C) and cooling (30 min of active cooling), the digests were filtered (qualitative filter paper, Grade 595 S+S, Whatman), quantitatively transferred and diluted with ultrapure water to a final volume of 50 mL prior to ICP-OES determination of elements. For river water, defined aliquots of river water were filtered (qualitative filter paper, Grade 595 S+S, Whatman), transferred to the test tubes and analysed directly according to the manufacturer’s recommendations. Water concentrations therefore refer to the operationally defined filtered fraction; metals bound to particles retained on the filter were not included in the water results.

### 2.3 Detection of trace metals (ICP-OES)

Analysis of element concentrations was carried out using an inductively coupled plasma optical emission spectrometer, Thermo iCAP 7600 Dual (Thermo Fisher Scientific, Waltham, MA, USA). The ICP-OES was operated at an RF plasma power of 1150 W, with coolant gas flow of 12 L/min, auxiliary gas flow of 0.50 L/min and nebuliser gas flow of 0.40 L/min. The nebuliser gas pressure was set to 120 kPa and the peristaltic pump speed to 50 rpm, under normal purge gas conditions. Calibration was performed using a Periodic table mix 1 for ICP (TraceCERT®, 33 elements, 10 mg/L in nitric acid; Sigma-Aldrich, product no. 92091). For ICP-OES measurement, the following wavelengths and instrumental limits of detection (LOD) and quantification (LOQ), expressed as concentrations in the measured solution, were as follows: Al 396.152 nm (LOD 0.002719 mg/L, LOQ 0.008239 mg/L), As 189.042 nm (LOD 0.004909 mg/L, LOQ 0.014876 mg/L), B 249.773 nm (LOD 0.000528 mg/L, LOQ 0.001600 mg/L), Ba 455.403 nm (LOD 0.000040 mg/L, LOQ 0.000121 mg/L), Be 313.042 nm (LOD 0.000050 mg/L, LOQ 0.000152 mg/L), Cd 228.802 nm (LOD 0.000263 mg/L, LOQ 0.000797 mg/L), Co 228.616 nm (LOD 0.000105 mg/L, LOQ 0.000318 mg/L), Cr 267.716 nm (LOD 0.000500 mg/L, LOQ 0.001515 mg/L), Cu 324.754 nm (LOD 0.000265 mg/L, LOQ 0.000803 mg/L), Fe 238.204 nm (LOD 0.000528 mg/L, LOQ 0.001600 mg/L), Mn 257.610 nm (LOD 0.000143 mg/L, LOQ 0.000433 mg/L), Ni 231.604 nm (LOD 0.001573 mg/L, LOQ 0.004767 mg/L), Pb 220.353 nm (LOD 0.002415 mg/L, LOQ 0.007318 mg/L), S 182.034 nm (LOD 0.029137 mg/L, LOQ 0.088294 mg/L), Si 251.611 nm (LOD 0.004594 mg/L, LOQ 0.013921 mg/L), Sr 421.552 nm (LOD 0.000043 mg/L, LOQ 0.000130 mg/L), V 292.402 nm (LOD 0.000278 mg/L, LOQ 0.000842 mg/L) and Zn 213.856 nm (LOD 0.000216 mg/L, LOQ 0.000655 mg/L). As water samples were measured without dilution, these limits apply directly to water; for sediments, the corresponding method limits are approximately 100-fold higher (0.5 g of sample digested and diluted to 50 mL). Analytical accuracy was verified with the certified reference materials ERM-CC141 (loam soil), ERM-CD281 (rye grass), ERM-CE278k (mussel tissue) and ERM-BB422 (fish muscle), all from the European Commission, Joint Research Centre (JRC), Directorate F – Health and Food (Geel, Belgium), which were analysed together with the samples; ERM-CC141 served as the matrix-matched material for sediments. Because neither the HNO₃/H₂O₂ microwave digestion used here nor the aqua regia extraction for which ERM-CC141 is certified decomposes silicates, the sediment concentrations represent pseudo-total (acid-extractable) contents. This fraction does not include elements bound within the silicate lattice, which are largely unavailable to biota, and it is commonly used in sediment risk assessment. Precision was evaluated from triplicate measurements, with relative standard deviations (RSDs) of 0.18–0.95% for all analysed elements.

### 2.4 Determination of total mercury by CV-AAS method

Total mercury (Hg) concentrations in river water and sediment were determined using a selective mercury analyzer AMA-254 (Altec, Prague, Czech Republic) based on cold vapour atomic absorption spectrometry (CV-AAS). The instrument operates at a wavelength of 253.65 nm, with thermal decomposition of the sample, amalgamation of elemental mercury on a gold trap and subsequent quantification of mercury vapour in an optical cell (Száková et al., 2004). For sediment, dried and homogenised samples were weighed into quartz sample boats (20 mg dry weight) and analysed directly without prior chemical digestion. Total mercury in water was measured from defined aliquots (100 µL), introduced directly into the liquid sample boats according to the instrument manufacturer’s instructions. The method detection limit for total mercury was 1.5 ng/kg (dry weight basis), and the limit of quantification was 4.5 ng/kg, in agreement with performance characteristics of the AMA-254 for environmental matrices. Analytical quality control was ensured by routine analysis of certified reference materials representative of soils/sediments (ERM-CC 141, European Commission, Joint Research Centre (JRC), Directorate F - Health and Food, Geel, Belgium), which were measured in each analytical batch alongside procedural blanks (six measurements). The recovery for the certified reference material was 97.01% of the certified value (Árvay et al., 2022; Demková et al., 2024)

### 2.5 Statistical evaluation

All analytical and visualization processes were performed using the programming environment R version 4.5.2 (R Core Team, 2025). In the initial phase, the dataset was subjected to a thorough cleansing process, with particular attention devoted to the management of values below the detection limit (LoD). Variables that exhibited an unacceptably high proportion of unquantifiable measurements were then excluded from the inferential analysis. Subsequently, we divided the remaining variables into two groups based on their relevance to the topic under investigation. Trace elements (Al, As, Cd, Cu, Fe, Hg, Mn, Ni, Pb, S, and Zn) directly related to our research were included in the main analytical matrix, and the other measured elements (B, Ba, Be, Co, Cr, Si, Sr, and V) are presented in the supplementary materials (SM1) in descriptive format.

Prior to the application of statistical tests, we systematically verified the assumptions of parametric testing for each element under study. We assessed the normality of the residual distribution using the Shapiro-Wilk test and evaluated the presence of homoscedasticity of variances among individual sampling sites using Levene’s test. Through the analysis of assumptions, we detected significant deviations from the normal distribution for most of the trace elements under investigation. For this reason, we chose a robust nonparametric approach for all spatial comparisons of concentrations across the river gradient.

We used the Kruskal-Wallis test to assess global differences in the concentrations of trace elements and heavy metals among sampling sites. In cases with a statistically significant global result, we followed up with Dunn’s post-hoc test, designed for detailed pairwise comparisons of specific locations. We corrected for the accumulation of Type I errors arising from multiple testing by applying the Holm-Bonferroni method. The level of statistical significance was set at α = 0.05 for all tests applied.

We quantified the environmental burden of sediments using two standardized metrics. We calculated the contamination factor (CF) as the ratio of the concentration of a specific metal in the evaluated sample to its average value at the reference site (L1). Subsequently, for each sample, we derived the pollution load index (PLI), defined as the geometric mean of the calculated contamination factors. We used software Labreador version 1.0.0 (Helczman, 2026) to calculate the CF and PLI.

We modelled the complex relationships between multiple variables and the spatial patterns of contamination using principal component analysis (PCA). We transformed the input data for the sedimentary subsystem using a decimal logarithm prior to principal component extraction. The purpose of this transformation was to stabilize the variance and appropriately scale the extreme values emitted by the primary source of pollution.

## 3. Results

### 3.1 Geochemical profile of the pollution source

In the initial phase of the analysis, we focused on a detailed quantification of contamination in the immediate area of the mine discharge. We defined the spatial gradient using five sampling points that map the condition of the stream upstream of the discharge (S4, S3), directly at the point of discharge (S0) of the mine water, and downstream of it (S1, S2). Surface water analysis revealed a localized increase in dissolved metal concentrations directly at the discharge point (Figure 2). We recorded Fe concentrations of 149.26 mg/L, Mn at 27.09 mg/L, and the presence of As at 0.29 mg/L. Outside this specific point, surface water showed a radically different result. Directly above the discharge point and immediately below it, Fe and As values fell below the limit of detection, representing a 100% proportion of non-quantifiable samples in these sections. Mn values beyond the direct source were also several orders of magnitude lower.

**Figure 2.**
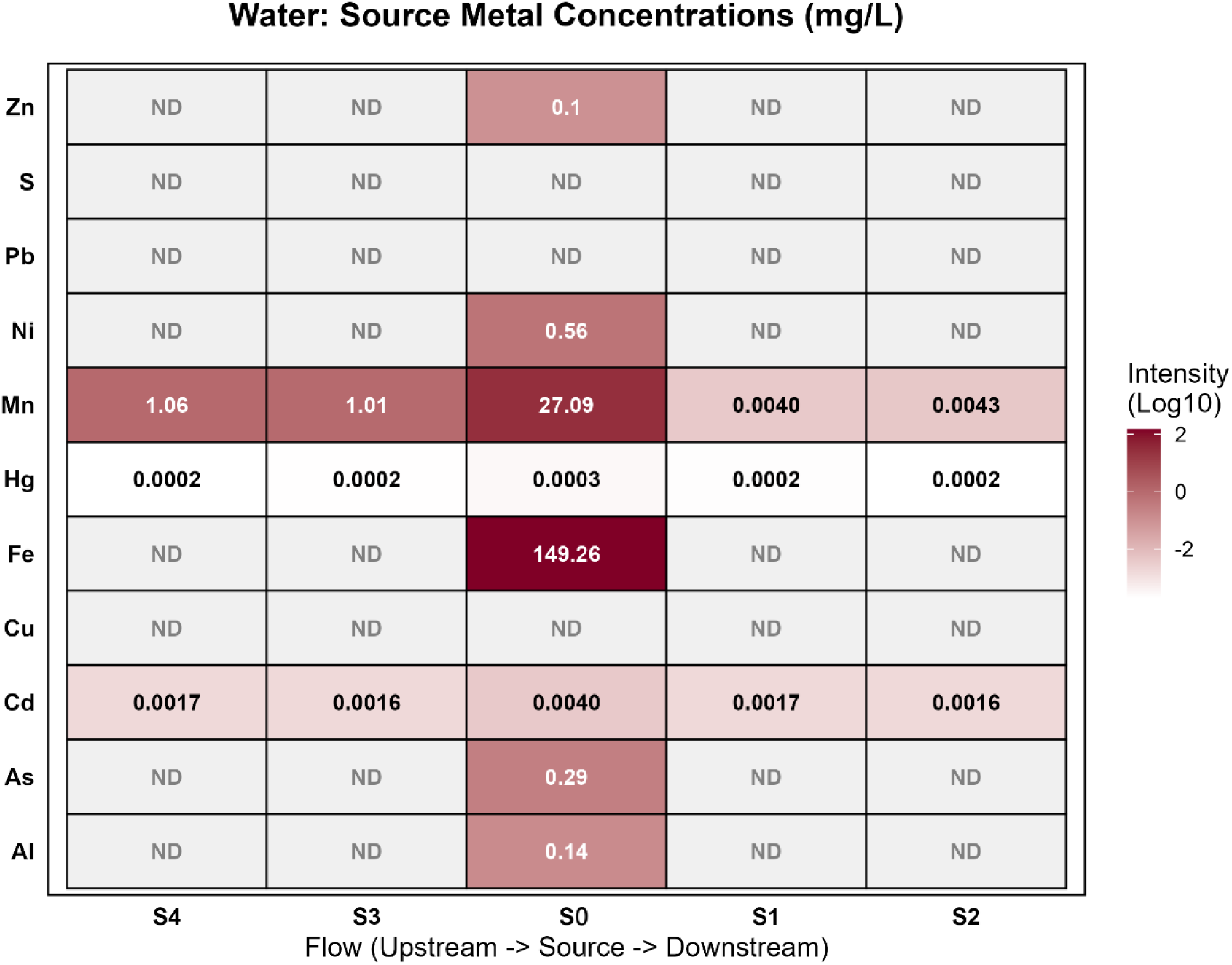
Relative intensity of heavy metal concentrations in surface water across the source micro-gradient. Note: Data represent mean raw concentrations (mg/L). ND indicates values below the limit of detection*; for sulphur (S), which was not determined in the water samples of the source micro-gradient, ND denotes “not determined”*.

In contrast to the water column, the sedimentary environment (Figure 3) exhibited more continuous spatial dynamics with a clearly recognizable accumulation of contaminants. We found that the deposition of solid particles massively traps precipitated metals. As in the sediment escalated from 83.78 mg/kg at a location 50 meters upstream of the discharge to 7920.28 mg/kg directly at the source. We identified a similar trajectory for sulphur, which increased from 195.94 mg/kg (S4) to 12 054.18 mg/kg at point S0. The elevated concentration of these elements persisted significantly even in sections immediately downstream of the pollution source. Cd showed the most pronounced local enrichment: it was not detected 50 m upstream of the discharge (S4) and reached only 0.12 mg/kg 20 m upstream (S3), whereas it rose to 38.95 mg/kg at the discharge point and remained high 20 m downstream (30.60 mg/kg), before decreasing to 5.00 mg/kg at S2. However, the spatial distribution was not uniform for all elements. In the case of Mn, we identified an inverse spatial pattern at the discharge site itself. While the concentration of this metal exceeded 1000 mg/kg at upstream locations, it dropped to 129.12 mg/kg directly at the source and subsequently rose again downstream. Fe maintained a relatively constant high concentration throughout the gradient, ranging from 10,450.49 to 14,088.53 mg/kg.

**Figure 3.**
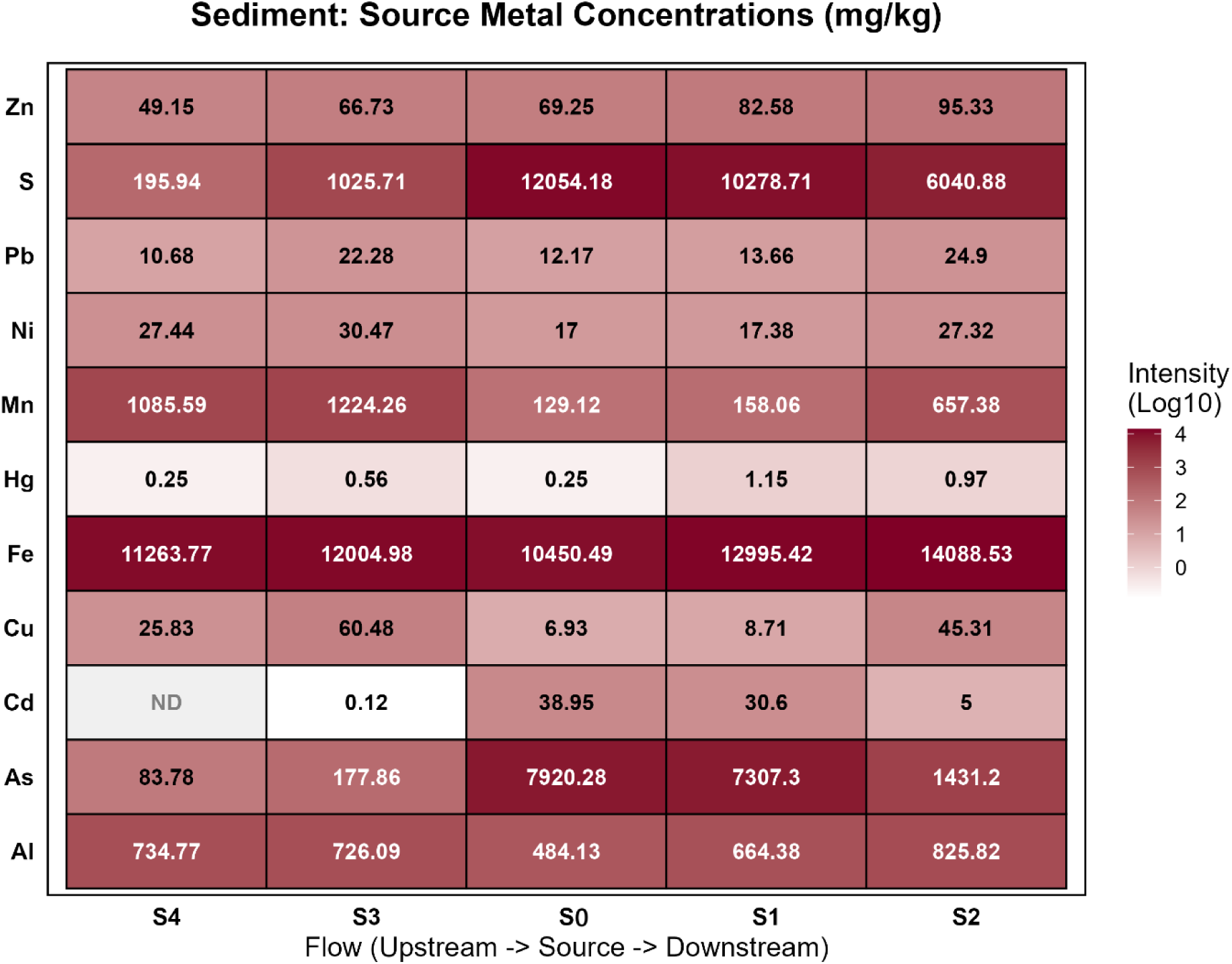
Relative intensity of heavy metal concentrations in sediment across the source micro-gradient. **Note:** Data represent mean raw concentrations (mg/kg).

#### River gradient - sediment

The spatial dispersion of contaminants along the river gradient exhibited heterogeneous patterns of decline (Figure 4). We applied the Kruskal-Wallis test and found statistically significant differences in As concentrations, H(3) = 23.242, p = 0.000036. A subsequent pairwise analysis using Dunn’s test with Holm-Bonferroni correction identified a primary increase in concentration between the reference site L1 and the first point downstream of the outfall L2, p = 0.002496. While As reached a level of 22.489 ± 3.233 mg/kg in the background of the river at L1, it increased to 47.189 ± 13.475 mg/kg at site L2. However, other sections of the river did not show a significant deviation from the reference values. Between L1 and L3, we recorded p = 0.502236, and between L1 and L4, p = 0.502236 as well. This indicates a relatively rapid downstream attenuation of As in the sediment. We observed very similar dynamics for Mn, H(3) = 22.602, p = 0.000049. The average Mn concentration at site L2 was 1436.676 ± 171.027 mg/kg and subsequently decreased significantly at site L3, p = 0.008538. We also observed this identical accumulation pattern downstream of the outfall for Hg, H(3) = 13.238, p = 0.004150. Its concentration increased from 0.227 ± 0.125 mg/kg at site L1 to 0.537 ± 0.222 mg/kg at site L2. We also identified an element with inverse spatial behaviour. Analysis of Pb revealed the presence of global variance, H(3) = 23.419, p = 0.000033. However, instead of the expected deposition at point L2, we measured the highest concentrations at the more distant site L3, where it reached 45.897 ± 83.274 mg/kg. A paired t-test confirmed a significant difference between L1 and L3, p = 0.010368. The marked heterogeneity of the data at this site represents a methodological limitation and implies the possible presence of an additional source of contamination.

**Figure 4.**
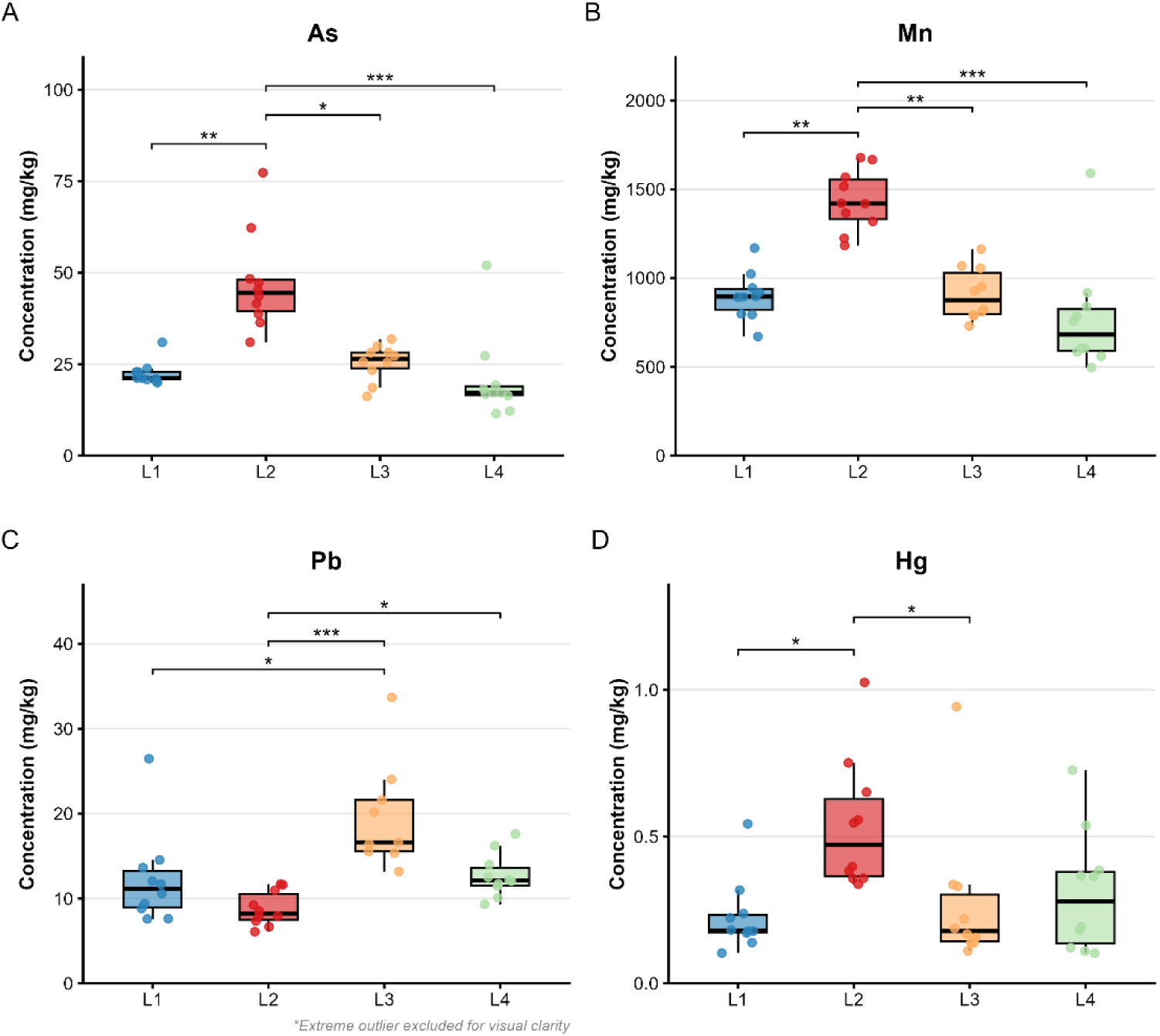
Distribution of heavy metal concentrations in sediment across sampling sites. **Note:** Boxes represent the interquartile range (IQR), and the horizontal line indicates the median. Significance levels: *** p < .001, p < .01, * p < .05, ns = not significant. Outliers are excluded from the visual range to improve clarity.

As shown in Figure 5, although the Fe testing detected a statistical difference between the distribution models, H(3) = 8.927, p = 0.030300, the absolute concentrations remained high throughout the entire monitored section. The difference between reference point L1, with a value of 10,962.849 ± 536.503 mg/kg, and impact zone L2, with a value of 11,488.925 ± 942.187 mg/kg, was not statistically significant. The only demonstrable decrease was observed between points L2 and L4, p = 0.021870. Ni values followed a similar pattern to those of As, S, and Hg. The average Ni value peaked at point L2 with a concentration of 31.947 ± 6.053 mg/kg and showed global statistical variance across the river, H(3) = 17.500, p = 0.000558. S values increased continuously downstream. A nonparametric test confirmed the presence of differences, H(3) = 18.028, p = 0.000434. Site L4 exhibited a significantly higher S level of 608.235 ± 279.660 mg/kg compared to zone L1, which had a concentration of 207.382 ± 120.263 mg/kg, p = 0.001150. We also observed a similar significant difference when compared to the other sites (L2, L3). A one-way analysis of variance for Al similarly demonstrated a linear increase in concentrations, F(3, 36) = 10.960, p = 0.000029. The Al content gradually increased from 554.910 ± 35.906 mg/kg at point L1 to 705.272 ± 63.100 mg/kg in zone L4. The remaining metals (Cu and Zn) exhibited less pronounced spatial dynamics. Cu values decreased slightly from L1 to L4 with global significance H(3) = 10.482, p = 0.014900. In contrast, Zn levels decreased at point L2 and subsequently increased at site L4, H(3) = 20.798, p = 0.000116.

**Figure 5.**
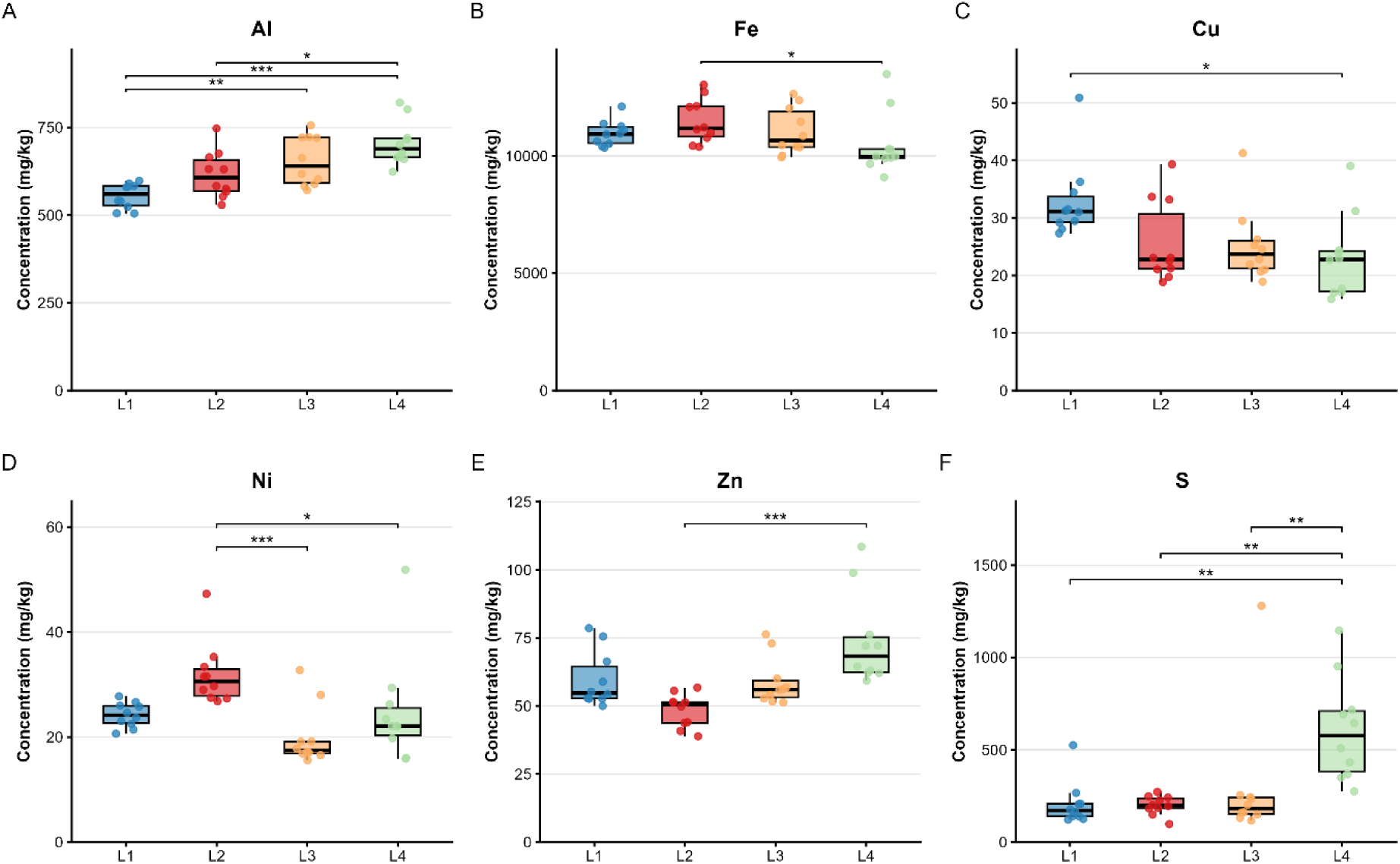
Distribution of associated trace elements in sediment across the river macro-gradient. **Note:** Boxes represent the interquartile range (IQR), and the horizontal line indicates the median. Significance levels: *** p < .001, p < .01, * p < .05, ns = not significant.

#### River gradient – water

Analysis of surface water along the river gradient (Figure 6) required the exclusion of almost all measured elements due to values below the limit of detection. For these reasons, we were able to quantify only S and Hg in the water. The distribution model of S in the aquatic environment showed statistically significant spatial variations, H(3) = 50.102, p < .001. At site L2, we observed a radical increase in S concentration to 35.697 ± 6.341 mg/L. Compared to the reference value of 6.372 ± 1.190 mg/L at point L1, this represents a significant change, p < .001. Increased distribution of S in the water persisted even at more distant sites, showing a downward trend. We continued to detect statistically significant differences between the reference site L1 and sites L3 (16.753 ± 1.397 mg/L) and L4 (17.503 ± 3.406 mg/L) (p = .003 and p < .001, respectively). A demonstrable decrease was recorded when comparing the zone of highest impact, L2, with the subsequent sections L3 (p < .001) and L4 (p = .003). For Hg, we identified global statistical variance H(3) = 11.005, p = .012. A subsequent post-hoc analysis confirmed the existence of a single significant difference, namely between the reference site L1 and the first contact point L2, p = .014; however, the absolute Hg concentrations at both sites were close to analytical zero.

**Figure 6.**
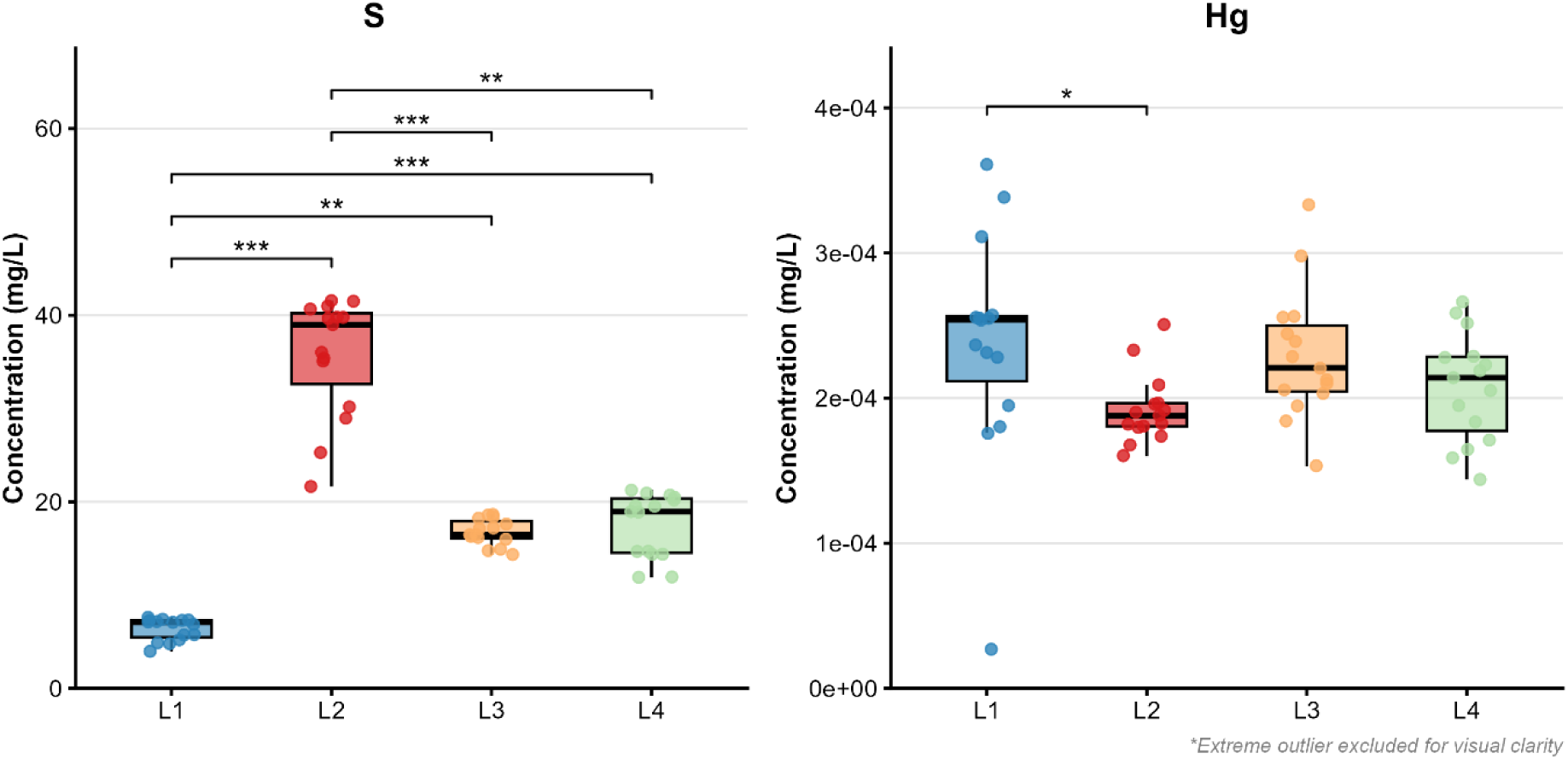
Spatial distribution of S and Hg in surface water across the river macro-gradient. **Note:** Boxes represent the interquartile range (IQR), and the horizontal line indicates the median. Significance levels: *** p < .001, p < .01, * p < .05, ns = not significant. Outliers are excluded from the visual range to improve clarity.

#### Principal component analysis

To identify the primary drivers of toxicity and assess the overall structural differences among individual sites, we applied principal component analysis (PCA). The first two principal components together explained 63.7% of the total spatial variance (PC1 = 36.4%; PC2 = 27.2%). Projection into a two-dimensional space revealed a clear separation of the mine discharge from the rest of the river gradient (Figure 7). Source (S0) represented a distinct outlier, whose position was determined primarily by strong vectors of As, S, and Hg. Despite the massive contamination directly at the discharge point, the PCA biplot showed surprisingly low spatial dispersion for the reference site L1 and the sites located downstream of the discharge (L2, L3, L4). These sampling points formed a relatively compact cluster with extensive overlap of their confidence ellipses. While zone L2 exhibited a slight shift in the direction of the Mn, Cu, and Ni vectors, the overall structure of the sedimentary matrix in areas L1 through L4 did not indicate the presence of radically different geochemical conditions in terms of the multivariate relationships of the monitored metals.

**Figure 7.**
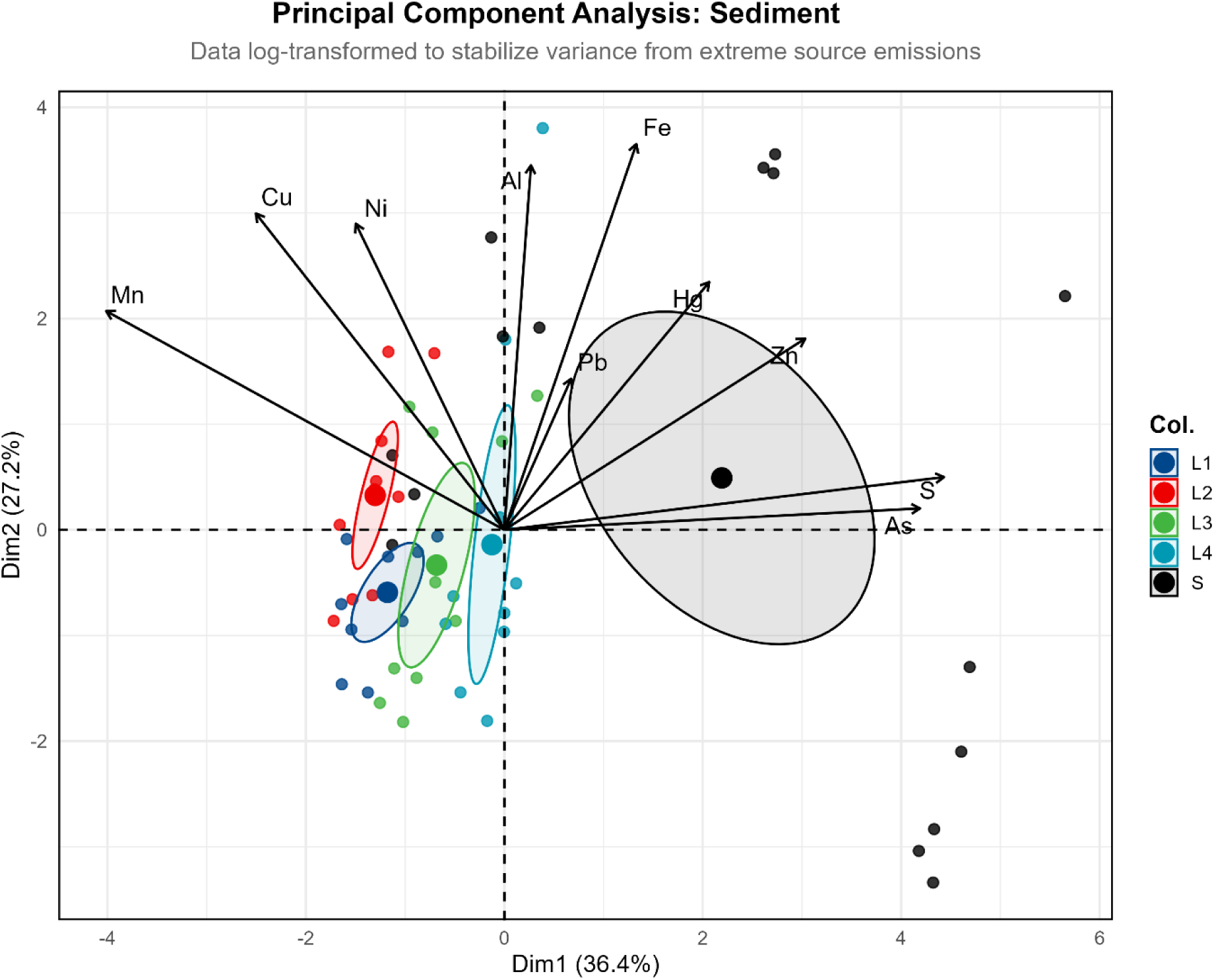
Principal Component Analysis (PCA) biplot of sediment profiles and inter-variable relationships. The model is based on Log10-transformed data to stabilize variance from extreme source emissions. PC1 (36.4%) and PC2 (27.2%) explain a total of 63.7% of the variance.

#### Contamination factor (CF) and pollution load index (PLI)

To verify the visual similarity of the sites (L1–L4) identified through PCA, we calculated the CF and, based on that, the PLI. This index aggregated the contributions of ten key elements into a single comparative metric, defined relative to the reference condition at site L1.

The PLI values indicate that all sites located downstream from the mine discharge are classified as polluted sections (PLI > 1). The highest index value was recorded in the immediate impact zone L2 at 1.16 ± 0.15 (Figure 8). The primary factors driving the index at this site were Hg (CF = 2.36), As (CF = 2.10), and Mn (CF = 1.60). More distant locations showed slightly lower but comparable values: L3 at 1.06 ± 0.20 and L4 at 1.10 ± 0.29. The most significant contributor in L3 was Pb (CF = 3.75), while in L4 it was the consistently increasing S (CF = 2.93). To statistically verify the differences between these three affected zones, we applied the nonparametric Kruskal-Wallis test, which did not confirm the existence of statistically significant differences in the total load of river sediment along the monitored gradient, H(2) = 2.805, p = 0.246.

**Figure 8.**
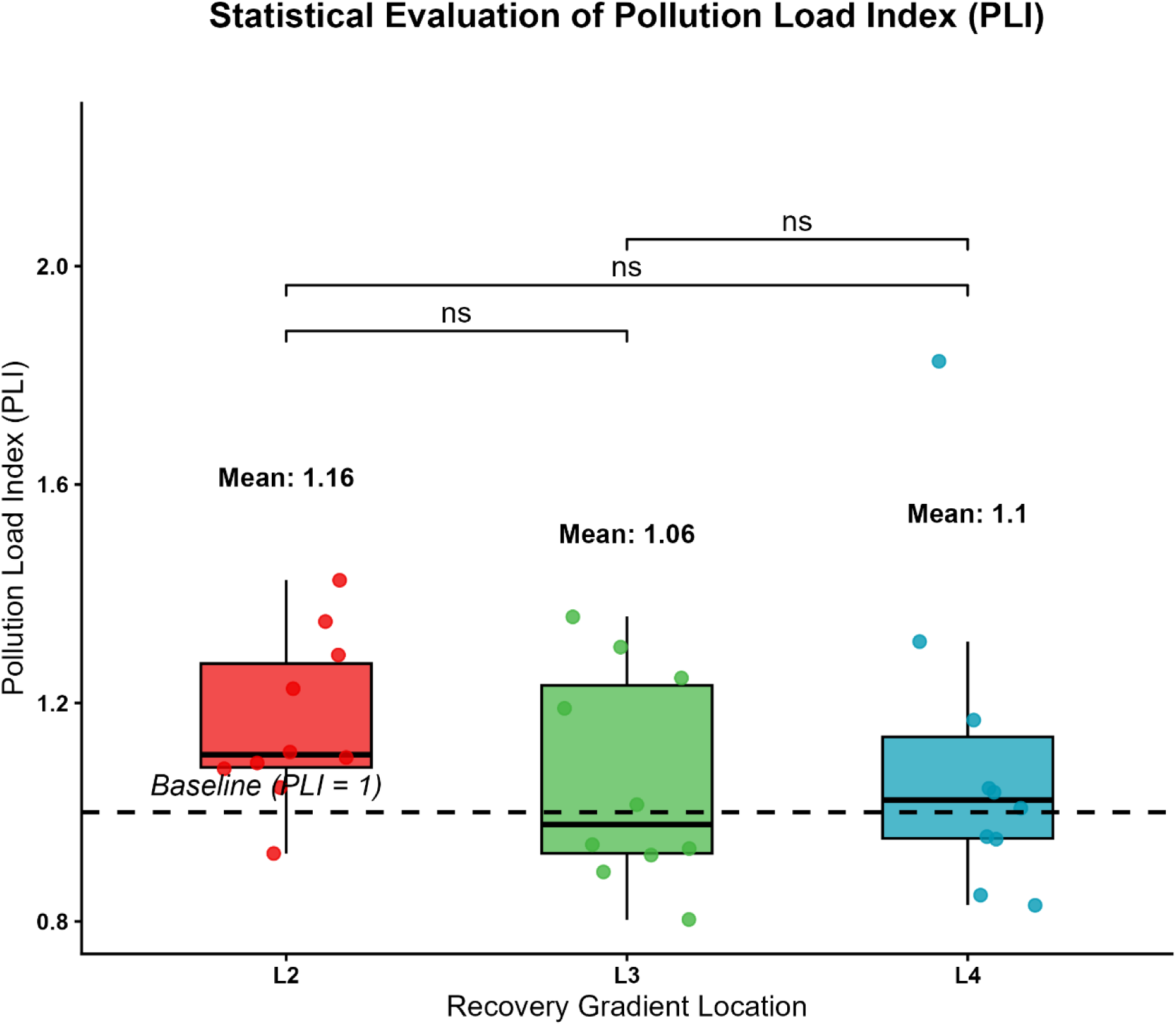
Assessment of the Pollution Load Index (PLI) across the river macro-gradient. **Note:** PLI represents the geometric mean of the calculated Contamination Factors (CF). Baseline is represented by reference site L1. Global significance (p = 0.246). No statistical difference in overall pollution burden between sites L2, L3, and L4. Values above 1 indicate a polluted status relative to the baseline.

## 4. Discussion

The results suggest that almost one and a half years after the environmental accident, the sedimentary component of the Slaná River remains a significant reservoir of contamination, even though most metals in the watercourse—excluding the area immediately downstream of the discharge point—have fallen below the limit of detection. A similar contrast between water and sediment has also been described in other areas affected by AMD, where there is a relatively rapid decline in the concentrations of dissolved metals in the water, but the sediments retain elevated contamination over the long term (Jennings et al., 2025; Stachnik et al., 2020). Fe(II) flowing from a mining source rapidly oxidizes to Fe(III), forming amorphous oxyhydroxides and hydrosulphates (such as ferrihydrite, schwertmannite, jarosite), which subsequently settle in the sediment and act as primary scavengers for oxyanionic pollutants such as As(V) and As (III) (Cheng et al., 2009; Gault et al., 2005; Morin et al., 2003), which may explain our extremely elevated As values (7920 mg/kg) directly at the source of pollution. An analogous dynamic was described by (Zhu et al., 2020) in a karst AMD system, where As and Fe in the water decreased from mg/L levels to trace values along the short flow, with almost the entire amount of pollutants remaining caught in the sediments. The same mechanism explains why Fe, Al, As, Ni and Zn, although quantified in the discharge water, were below the LOD in the filtered river water already 20 m downstream of the source: once the mine water mixes with the receiving river, these elements are rapidly transferred to Fe/Al oxyhydroxide and hydroxysulphate particles, which were retained on the filter but are reflected in the strong enrichment of the sediment (e.g., 7307 mg/kg As 20 m downstream). However, Jung et al., (2025) document that As contamination in similar mining-impacted systems can migrate as far as 50 km downstream despite sorption and coprecipitation processes. The concentrations of Fe (149 mg/L), Mn (27 mg/L), and As (0.29 mg/L) directly at the discharge point are consistent with the geochemistry of siderite-pyrite-arsenopyrite deposits and correspond to the profile described for the locality of Nižná Slaná itself. Mačingová et al. (2024) describe alternative approaches to removing metals from the same mine stream in their work, mentioning high concentrations of Fe and Mn in this mine water. Ulytskyi et al. (2025), in a study directly devoted to the ecological disaster on the Slaná River, confirm elevated temperature, reduced dissolved oxygen content, lowered pH, and high concentrations of sulphates, Fe, As, Co, and Zn in the discharge zone. Precisely the lowered pH in this area provides a possible physicochemical explanation for the inverse spatial pattern of Mn that we observed (higher concentrations in the sediment above the discharge point than in the discharge itself). Mn(II) is significantly slower in terms of oxidation kinetics and requires a higher pH (typically > 8) for effective abiotic precipitation (Li et al., 2019; Neculita C Rosa, 2019). In the slightly acidic to neutral zone, Mn remains predominantly in the dissolved form—which is why we observe a depression (129 mg/kg) in the discharge sediment despite elevated concentrations in the water.

Our data distinguish two distinct types of pollutant attenuation along the river gradient in sediments. The first group (As, Hg, Mn, and Ni) is characterized by acute deposition in the impact zone (L2) and subsequent stabilization toward reference values. This decline is again consistent with the mechanism of Fe-oxyhydroxide coprecipitation and sedimentation of coarser aggregates in the immediate neutralization zone (Lezar C Akkal, 2026; Tabelin et al., 2020). The second group (S and Al) is characterized by the opposite, cumulative pattern, with a maximum at the most distant location, L4. While iron precipitates immediately, Al requires a higher pH (>5.0) for hydrolysis and precipitation. The gradual neutralization of the river toward L4 triggers the precipitation of Al-hydrosulphate phases (e.g., basaluminite), which explains the simultaneous uptake of both aluminium and sulphates from the water column into the sediment, supplemented by potential microbial sulphate reduction in distal sections (Fernando et al., 2018; Galhardi C Bonotto, 2016; Xia et al., 2017).

One of the most important, yet easily overlooked, results of our PCA is the relative compactness of the L1–L4 cluster and the significant overlapping of their confidence ellipses. This finding supports a critical interpretive limitation. The reference site L1 may not represent a truly “natural” geochemical background, but rather a regional background already historically burdened by long-term mining activity. Fazekašová C Fazekaš (2020) in a study of soils in the Nižná Slaná emission field, document that the concentrations of Hg, Cd, Cr, Cu, As, Fe, Mn, and Mg are above the toxicological threshold in the wider area. Similar findings in the surrounding soil in these areas are also reported by Demková et al., (2020), who calculated elevated PLI values in nearly all observations. The PCA-detected homogeneity of L1–L4 may not signal effective dilution of the source but merely reflects a “shifted baseline”—the baseline level of contamination against which any local gradient may appear attenuated.

To evaluate the toxicological relevance of the measured concentrations, we compared them with the consensus-based sediment quality guidelines for freshwater ecosystems (MacDonald et al., 2000), which define a threshold effect concentration (TEC), below which adverse effects on benthic organisms are unlikely, and a probable effect concentration (PEC), above which such effects are expected to occur frequently. Arsenic was the most critical element: at the discharge point (7920 mg/kg) and 20 m downstream (7307 mg/kg), its concentrations exceeded the PEC (33 mg/kg) more than 200 -fold, and the mean value at the first downstream river site L2 (47.2 mg/kg) remained above the PEC. Cadmium, which was not detected 50 m upstream of the discharge, reached 38.95 mg/kg at the discharge and 30.60 mg/kg 20 m downstream, i.e. approximately six- to eightfold the PEC (4.98 mg/kg), and decreased to the PEC level 50 m downstream (5.00 mg/kg). The mine discharge thus represents a highly localized but intense source of Cd, whose sedimentary footprint is shorter than that of As. Hg exceeded the PEC (1.06 mg/kg) 20 m downstream of the discharge (1.15 mg/kg), and the mean concentrations of Hg and Ni at L2 exceeded their respective TECs. Notably, As concentrations at both upstream micro-gradient points (83.8 and 177.9 mg/kg) and the mean As and Hg concentrations at the reference site L1 (22.5 and 0.23 mg/kg) also exceeded the TEC or even the PEC, which supports the interpretation of a historically elevated regional baseline discussed above. Because the HNO₃/H₂O₂ digestion yields pseudo-total contents, these exceedances should be regarded as conservative estimates.

PLI values between 1.06 and 1.16 in all affected zones formally confirm classification as contaminated sections, but they are relatively low compared to other mining-impacted ecosystems. Sabijon et al. (2025) report a PLI > 1.5 in the post-mining Taft River (Philippines); Pedrozo-Acuña et al. (2025) describe a PLI > 3 in the deposition zones of the Sonora River. Our values fall within the range reported for the mining-affected Emet River basin in Turkey (PLI 0.45–1.20), where As, Cd and Pb were likewise identified as the elements of greatest concern in sediments (Tokatli et al., 2021). Our moderate PLI values may have been suppressed by the previously discussed effect of the shifted L1 reference. Our single-campaign data from one and a half years after the accident suggestthat, apart from sulphur, the filtered water column has largely returned to near-background levels, but the sediment remains a persistent latent source of contamination. The stability of Fe/Mn-oxyhydroxide-associated metals is strictly dependent on redox and pH parameters. A decrease in redox potential due to anoxia or acidification can release As, Pb, or other heavy metals back into the water column (Hao et al., 2021; Madadi et al., 2021; Yao et al., 2024). Specifically for As, Tabelin et al., (2020) and Zhang et al., (2022) show that As mobility exhibits a V-shaped dependence on acidity with a minimum around neutral conditions, while deviations in acidity in both directions increase leachability. Experience from comparable mining disasters confirms that the recovery trajectory is long and often incompletely successful. Baos et al. (2022) observed persistent contamination by Cd, Pb, and As in the Guadiamar River (the 1998 Aznalcóllar accident) even 8 years after the accident, along with occasional rebounds; Soares et al. (2025) in a systematic review of the consequences of the Fundão dam collapse (Mariana, Brazil, 2015), describe lasting physical and chemical changes in the Doce River as late as 7 years after the event.

Study limitations: Several limitations should be considered when interpreting our results. First, the data derive from a single sampling campaign conducted in September 2023. Seasonal changes in discharge, precipitation and temperature can substantially alter the partitioning of metals between water and sediment, as well as the resuspension and downstream transport of contaminated sediment (Jennings et al., 2025; Jung et al., 2025). Our findings therefore represent a snapshot rather than a description of the long-term trajectory of the river, and statements regarding stabilization or recovery should be regarded as preliminary. Second, the water samples represent the operationally defined filtered fraction; metals associated with suspended particles were not captured in the water phase, and all elements except S and Hg had to be excluded from the analysis of the river gradient because most values were below the LOD. Third, sediment concentrations were determined after HNO₃/H₂O₂ digestion and represent pseudo-total contents, which may underestimate the total concentrations of elements bound in the silicate matrix. Finally, the reference site L1 is located in a historically mining-affected area and probably represents a regional rather than a natural background, which may have lowered the calculated CF and PLI values. Repeated sampling across hydrological seasons, including suspended particulate matter, together with the ongoing analysis of fish from the same sites, is needed to verify the persistence of the observed patterns.

## 5. Conclusion

One and a half years after the mine-water spill, the water column of the Slaná River showed only a local contamination signal: apart from the discharge itself, most metals in filtered water were below the detection limit, and only sulphur remained elevated downstream. The sediments, however, still retain the contamination. Arsenic and cadmium at the discharge, and arsenic at the first downstream site, exceeded the probable effect concentrations of freshwater sediment quality guidelines, and even the upstream reference reach exceeded threshold levels for As and Hg, reflecting a historically elevated regional background. Because the downstream decline of the primary mining contaminants (As, Hg, Mn, Ni) is offset by the accumulation of S and Al, the overall pollution load remains uniform along the studied reach. The main implication of this study is that improved water quality alone does not indicate recovery of the Slaná River: the sediment remains a persistent and potentially remobilisable reservoir of toxic elements, which should be considered in further monitoring and in interpreting effects on fish. As these conclusions rest on a single sampling campaign, repeated seasonal sampling is required to confirm them.

## Funding

This work was supported by The Ministry of Education, Research, Development and Youth of the Slovak Republic under the project VEGA 1/0571/23 and by the Slovak Research and Development Agency under the contract No. APVV-21-0168.

## Author Contributions (CRediT)

**Marek Helczman:** Conceptualization, Methodology, Software, Validation, Formal analysis, Data Curation, Writing - Original Draft, Visualization; **Marian Tomka:** Methodology, Validation, Formal analysis, Investigation, Writing - Review C Editing; **Anton Kovacik:** Conceptualization, Resources, Writing - Original Draft, Writing - Review C Editing, Visualization, Supervision, Project administration, Funding acquisition

## Consent to Participate

All authors confirm to participate in this study.

## Conflicts of Interest

The authors declare no conflict of interest.

## Data availability

Data will be made available from corresponding author on reasonable request.

## Generative AI statement

The authors state the use of generative artificial intelligence tools in the preparation of this manuscript. Specifically, Perplexity Pro (© 2026 Perplexity AI) and DeepL Translator (DeepL SE) were utilized to assist with language refinement, including translation and grammar checking, with the aim of improving clarity and academic quality. All outputs generated by these tools were carefully reviewed and edited by the authors, who retain full responsibility for the final content of the publication.

## Notes

### Competing Interest Statement

The authors have declared no competing interest.

### Summary of Updates

The abstract, Figure 1, and conclusions have been revised. The author list has also been updated to remove Dr. L. Harangozo, who has left academia and requested to no longer be included as a co-author.

