## Supplemental Table 1 and Table 2 for "Assessment of trace metals and arsenic contamination in water and sediments in a former mining area of the river basin after an ecological disaster, Slaná River, Slovakia"

**SM1 Table 1** Average values of elements not included in further analysis at the source of contamination (results are presented as mean ± SD)

| Matrix | Location | B | Ba | Be | Co | Cr | Si | Sr | V |
|---|---|---|---|---|---|---|---|---|---|
| Sediment | S | 262.350 ± 15.795 | 7.647 ± 1.102 | 1.664 ± 0.151 | 0.378 ± 0.086 | 2.636 ± 0.310 | 88.909 ± 6.128 | 3.520 ± 0.435 | 2.639 ± 0.245 |
| Sediment | S1 | 260.456 ± 21.273 | 10.351 ± 0.569 | 1.888 ± 0.097 | 0.753 ± 0.039 | 3.104 ± 0.167 | 192.724 ± 75.389 | 5.063 ± 1.042 | 2.824 ± 0.070 |
| Sediment | S2 | 69.784 ± 2.584 | 95.224 ± 1.601 | 0.616 ± 0.027 | 10.725 ± 0.418 | 20.372 ± 2.163 | 169.538 ± 27.874 | 15.511 ± 0.693 | 14.825 ± 0.512 |
| Sediment | S3 | 41.332 ± 3.133 | 93.491 ± 10.585 | 0.393 ± 0.025 | 9.160 ± 0.644 | 22.817 ± 4.123 | 225.190 ± 51.129 | 18.495 ± 2.400 | 15.344 ± 1.965 |
| Sediment | S4 | 29.666 ± 0.916 | 59.494 ± 3.537 | 0.277 ± 0.044 | 8.648 ± 0.506 | 23.150 ± 3.196 | 183.410 ± 3.031 | 13.651 ± 1.750 | 12.354 ± 1.136 |
| Water | S | ND | 0.061 ± 0.004 | ND | ND | ND | ND | ND | ND |

| Matrix | Location | B | Ba | Be | Co | Cr | Si | Sr | V |
|---|---|---|---|---|---|---|---|---|---|
| Water | S1 | ND | 0.039 ± 0.003 | ND | ND | ND | ND | ND | ND |
| Water | S2 | ND | 0.044 ± 0.000 | ND | ND | ND | ND | ND | ND |
| Water | S3 | ND | 0.030 ± 0.003 | ND | ND | ND | ND | ND | ND |
| Water | S4 | ND | 0.030 ± 0.000 | ND | ND | ND | ND | ND | ND |

**SM1 Table 2** Average values of elements not included in further analysis in in the surveyed locations (results are presented as mean ± SD)

| **Matrix** | **Location** | **B** | **Ba** | **Be** | **Co** | **Cr** | **Si** | **Sr** | **V** |
|---|---|---|---|---|---|---|---|---|---|
| Sediment | L1 | 26.630 ± 2.640 | 87.273 ± 13.633 | 0.295 ± 0.066 | 8.693 ± 0.889 | 26.499 ± 4.439 | 217.219 ± 23.143 | 17.181 ± 2.823 | 14.377 ± 2.531 |
| Sediment | L2 | 31.212 ± 4.647 | 92.162 ± 22.063 | 0.301 ± 0.041 | 10.210 ± 1.247 | 25.422 ± 9.974 | 242.472 ± 21.435 | 17.169 ± 2.371 | 10.521 ± 2.070 |
| Sediment | L3 | 27.817 ± 3.627 | 99.327 ± 20.635 | 0.255 ± 0.043 | 9.935 ± 1.517 | 17.402 ± 1.848 | 196.783 ± 19.973 | 19.195 ± 2.044 | 13.528 ± 1.526 |
| Sediment | L4 | 25.056 ± 5.848 | 100.703 ± 26.216 | 0.317 ± 0.124 | 10.556 ± 2.844 | 20.479 ± 3.462 | 172.413 ± 33.499 | 25.011 ± 4.426 | 21.272 ± 4.492 |
| Water | L1 | ND | 0.036 ± 0.004 | ND | ND | ND | 5.066 ± 0.633 | 0.110 ± 0.009 | ND |
| Water | L2 | ND | 0.009 ± 0.004 | ND | ND | ND | 4.746 ± 0.519 | 0.092 ± 0.017 | ND |

| Matrix | Location | B | Ba | Be | Co | Cr | Si | Sr | V |
|---|---|---|---|---|---|---|---|---|---|
| Water | L3 | ND | 0.022 ± 0.003 | ND | ND | ND | 6.057 ± 0.146 | 0.147 ± 0.009 | ND |
| Water | L4 | ND | 0.020 ± 0.002 | ND | ND | ND | 5.986 ± 0.436 | 0.149 ± 0.023 | ND |
